# Engineering Persistent and Rewritable Genetic Memory in Bacteria through Multiscale Plasmid Competition

**DOI:** 10.64898/2026.09.18.752696

**Authors:** Iacopo Ruolo, Eric Lu, Domitilla Del Vecchio

## Abstract

Genetic memory enables bacteria to record transient stimuli as persistent genetic states, providing a foundation for cellular information storage, sensing, and computation. Recombinase-based memories encode discrete states through rewritable DNA rearrangements, but these states can continue to evolve after stimulus removal as cells proliferate. Here, we show that the long-term stability of reversible recombinase-based memory can be engineered by controlling post-switching dynamics. We identify a temporal hierarchy in which heterogeneous states arising from imperfect recombinase orthogonality are first resolved through intracellular competition between incompatible plasmids and subsequently reshaped by fitness differences between cells carrying alternative states. Reducing DNA copy number accelerates intracellular state resolution and decreases expression-associated fitness costs, while tuning state-specific gene expression balances the relative fitness of alternative memory states. Using these principles, we engineer a reversible memory device that maintains its two states for up to two weeks while retaining rewritability after prolonged maintenance. Our results establish a framework for engineering persistent and rewritable genetic memory through control of intracellular competition and population-level fitness.

## Introduction

Synthetic gene circuits that store information in bacterial cells are fundamental components of cellular computation ^1–6^. By converting transient inputs into persistent molecular states, genetic memory enables cells to record past events and make decisions based on their history, with applications ranging from biological computation and diagnostics to therapeutics and biocontainment ^7–11^. Despite extensive efforts to engineer increasingly sophisticated memory devices, maintaining a defined state over extended periods of time remains challenging. In bacterial systems, memory can decay over timescales of days after removal of the input stimulus, limiting applications that require durable information storage ^12,13^. Thus, establishing what determines the long-term fate of synthetic genetic memory in a growing bacterial population could provide valuable information for engineering long-term, rewritable memory devices.

At the molecular level, genetic memory in bacterial cells can be implemented through regulatory feedback, like epigenetic mechanisms, or DNA rearrangements that generate persistent genetic states ^2–6,14,15^. Recombinase-based systems are particularly attractive because programmable DNA inversions can encode discrete states and, in reversible configurations, allow information to be written, erased, and rewritten in response to transient signals ^10,12,16–22^. Yet, reversible DNA inversion can introduce genetic heterogeneity during switching. In multicopy plasmid-based systems, incomplete or unintended recombinase activity, including leaky expression and limited recombinase orthogonality, can prevent all plasmid copies from adopting the same configuration, thereby reducing switching efficiency and generating heterogeneous genetic states. Consequently, cells may retain mixtures of plasmid variants with opposite promoter orientations, corresponding to alternative transcriptional states ^10,14,16,17^. The state observed at the cellular level may therefore depend not only on the efficiency of the recombination reaction, but also on the subsequent dynamics of the competing genetic states generated during switching.

Competition between genetic elements is a fundamental property of plasmid biology and can operate at multiple levels. Plasmid variants sharing replication or partitioning systems can compete within individual cells ^23–28^, while cells carrying different plasmid compositions can compete with one another at the population level ^12,29^. Plasmids belonging to the same incompatibility group, for example, cannot generally be stably maintained together because they interfere with one another’s replication or partitioning, ultimately favoring the loss of one variant ^29–35^. At the population level, plasmid carriage and expression can alter host fitness, causing cells carrying different plasmid variants to proliferate at different rates. Consequently, the fate of a plasmid genotype can be shaped by both intracellular competition and intercellular selection ^12,29,36–38^.

Recent studies have emphasized the importance of distinguishing between these two levels of competition. Bedhomme et al. showed that plasmids can experience competition within individual cells while their host clones simultaneously compete at the population level, demonstrating that these processes can jointly shape plasmid evolutionary dynamics ^38^. More recently, Rossine et al. showed that intracellular competition between plasmids can make a transient but consequential contribution to plasmid population dynamics and interact with fitness-based selection acting between cells ^29^. These studies establish that intracellular and population-level competition represent distinct processes that operate on different timescales and can favor different genetic outcomes. They therefore provide a framework for considering genetic memory not simply as a molecular state encoded by a circuit, but as a state whose persistence can be shaped by interactions occurring both within cells and across a growing population.

The relevance of these processes to genetic memory is further supported by studies showing that cellular fitness can influence the persistence of stored genetic states. Kalvapalle et al. demonstrated that fluorescent output associated with a recombinase-recorded state imposes a fitness cost that contributed to progressive memory loss over several days. Notably, removing the fluorescent output, by reducing the burden associated with the stored state, substantially improved memory persistence ^13^. This work showed that cellular burden and fitness can contribute to long-term memory stability. Together with studies of plasmid competition, these findings suggest that a memory state emerges from both plasmid competition within individual cells and by the differential expansion of cells carrying alternative states.

Rather than viewing these processes solely as limitations to be eliminated, we consider them as engineering variables for long-term memory. Characterizing these processes within a single reversible memory system could thus reveal how intracellular genetic competition, plasmid copy number, and state-dependent cellular fitness contribute to memory dynamics and how these parameters can be deliberately tuned to extend memory persistence.

Here, we combine experiments with stochastic computational modeling to follow the fate of competing plasmid states from the immediate switching through long-term population growth. We first characterize the resolution of heterogeneous plasmid states within individual cells and determine how plasmid copy number influences this process. We then examine the longer-term dynamics of the resulting memory states and quantify the contribution of differential cellular fitness to population-level state persistence. Finally, we use plasmid copy number and state-specific resource loading to reshape the relative fitness of the two memory states while retaining their persistence. This strategy extends the memory duration, with the optimized switch maintaining its programmed state for up to two weeks. Together, our results demonstrate that the long-term persistence of reversible genetic memory can be engineered by modulating the intracellular and population-level processes that govern the fate of competing genetic states.

## Results

### Design of a reversible recombinase-based toggle switch

To investigate the determinants of long-term memory persistence in a reversible recombinase-based genetic system, we engineered a two-plasmid toggle switch based on a previously described design ^16^ (Figs. 1A and S1A).

**Fig. 1.**
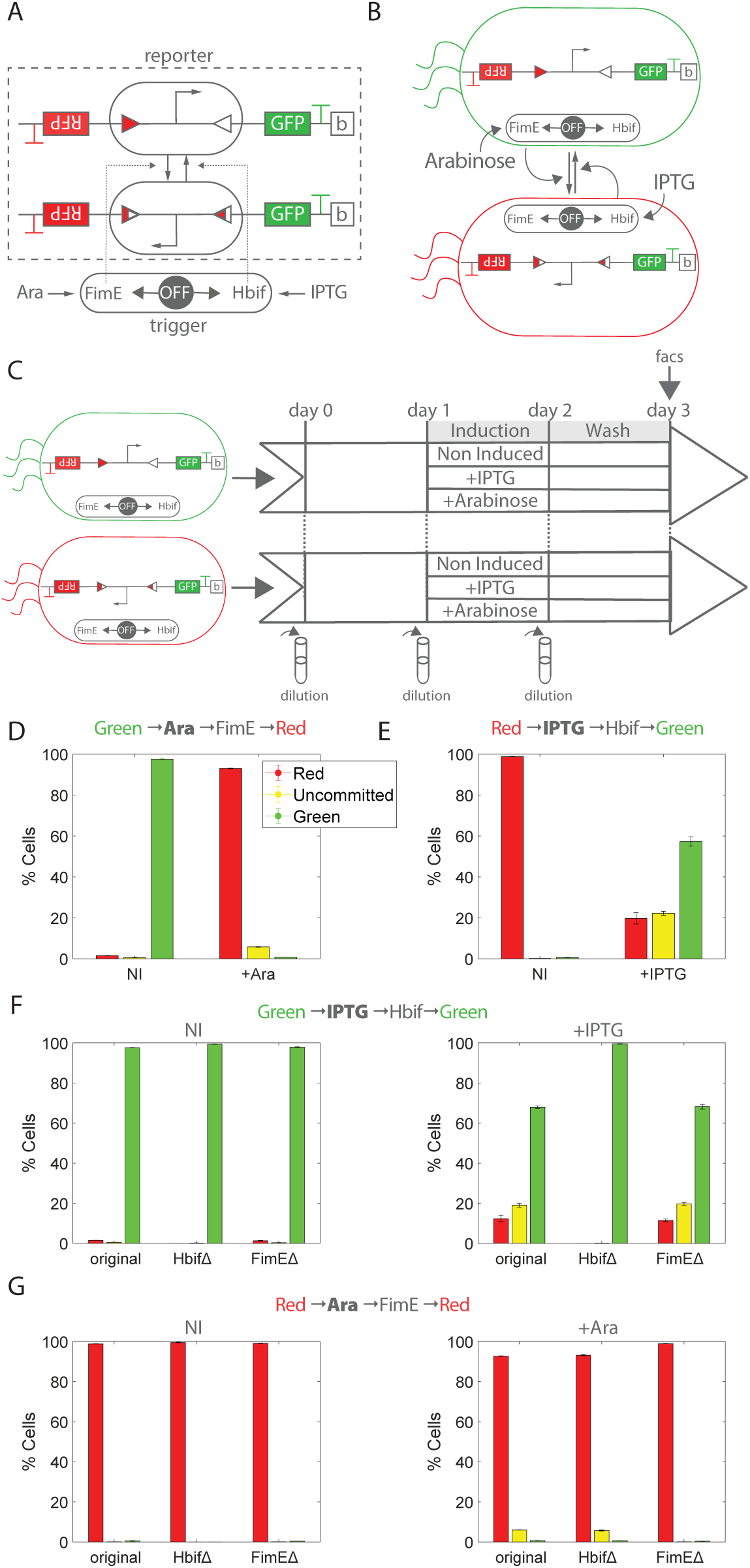
Recombinase-based bacterial switch: assessment of recombinase orthogonality. (A) Schematic representation of the two-plasmid recombinase-based memory switch, comprising the complementary irreversible recombinases FimE and HbiF. (B) Cells initialized in the Green-On state can be switched to the Red-On state by induction with arabinose. Conversely, cells initialized in the Red-On state can be switched to the Green-On state by induction with IPTG. The populations are switched according to the “Switching between states” protocol described in Methods. (C) Schematic representation of the cell-switching treatment protocol. (D) Cells initialized in the Green-On state and switched to the Red-On state on day 3, together with the non-induced control (NI). (E) Cells initialized in the Red-On state and switched to the Green-On state on day 3, together with the non-induced control (NI). (F) Cells initialized in the Green-On state and induced with IPTG, together with the non-induced control (NI), for constructs co-transformed with three trigger variants: one enabling inducible production of both FimE and HbiF (original), one enabling production of FimE only (HbiFΔ), and one enabling production of HbiF only (FimEΔ). (G) Cells initialized in the Red-On state and induced with arabinose, together with the non-induced control (NI), for constructs co-transformed with three trigger variants: one enabling inducible production of both FimE and HbiF (Total), one enabling production of FimE only (HbiFΔ), and one enabling production of HbiF only (FimEΔ). Details on how the percentage of green, red, and uncommitted cells are evaluated are reported in Fig. S1.

The input module (trigger) contains IPTG- and L-arabinose-inducible promoters driving the expression of HbiF and FimE, respectively. The output module (reporter) contains recombinase recognition sites flanking a constitutive promoter (Ptrc*) controlling RFP or GFP expression. Site-specific DNA inversion between the inverted repeats (IRL/IRR) reconfigures the recombination sites, reversing promoter orientation and thereby toggling expression between RFP and GFP. The reporter cassette was engineered on a backbone carrying a five-copy pSC101 origin of replication ^39^.

The system was designed to establish two alternative transcriptional states that could be maintained in the absence of induction and reversibly switched following transient activation of the appropriate recombinase. Thus, a successful memory transition requires not only efficient recombination, but also subsequent maintenance of the newly established state after removal of the inducing signal.

To characterize the establishment of the two memory states, we generated two bacterial populations carrying the same trigger plasmid but a reporter plasmid initialized in either the GFP-producing or RFP-producing state (Fig. 1B). GFP-On cells were induced with L-arabinose to promote switching to the RFP-On state, whereas RFP-On cells were induced with IPTG to promote switching to the GFP-On state (Fig. 1C).

Switching from the GFP-On to the RFP-On state reached approximately 95% efficiency (Fig. 1D), whereas switching from the RFP-On to the GFP-On state reached approximately 60% efficiency (Fig. 1E) two days after induction on day 1. Thus, although both transitions were functional, switching generated asymmetric populations containing different proportions of cells that had not yet adopted the desired state.

We first investigated whether incomplete switching could be attributed to basal activity of the trigger. Focusing on the non-induced (NI) controls in Fig. 1D and E, both populations remained stable over time in the absence of induction. These results indicate that basal expression of the trigger was not sufficient to induce substantial state switching and therefore exclude trigger leakiness as the primary cause of the incomplete transitions.

We next investigated whether the incomplete transitions resulted from insufficient recombinase orthogonality, unintended effects of the inducing molecule, or transcriptional readthrough ^2,14,16,17,40,41^. To specifically assess the orthogonality of HbiF, GFP-On cells were induced with IPTG, which activates HbiF but should not induce a transition from the GFP-On state (Fig. 1F). When cells carrying the complete trigger (original) were exposed to IPTG, a substantial fraction of the population was displaced from the GFP-On state by day 3, when instead no transition should have been observed. This result indicated that activation of HbiF could perturb the GFP-On state.

To distinguish between possible mechanisms underlying this response, we generated two trigger variants lacking either HbiF (HbiFΔ) or FimE (FimEΔ). Induction of GFP-On cells carrying the HbiFΔ trigger produced no detectable change in population state, excluding a direct effect of IPTG and indicating that the inducing molecule itself was not responsible for the observed phenotype. The absence of a response in this condition also excludes transcriptional readthrough through FimE as the cause of the observed switching. In contrast, induction of cells carrying the FimEΔ trigger, in which HbiF was the only inducible recombinase, reproduced the state displacement observed with the complete trigger. Together, these results demonstrate that HbiF is not fully orthogonal to the GFP-On state and can induce unintended switching upon activation.

We obtained analogous results when examining the reciprocal transition by inducing RFP-On cells with Larabinose (Fig. 1G). Induction of cells carrying the FimEΔ trigger, in which HbiF was absent, produced no detectable change in the population, excluding an effect of arabinose itself. Similarly, the absence of a response in this condition excludes transcriptional readthrough through HbiF as the cause of the phenotype. In contrast, induction of cells carrying the HbiFΔ trigger, in which FimE was the only inducible recombinase, resulted in a detectable displacement of the population from the RFP-On state, confirming that FimE also exhibits incomplete orthogonality, although to a less extent.

Taken together, these findings demonstrate that the two recombinases are not orthogonal in the engineered circuit. Activation of either recombinase can therefore perturb the opposite memory state, generating heterogeneous populations following a switching event. Thus, incomplete recombinase orthogonality represents an important determinant of how the initial distribution of genetic states is established following induction, providing a potential source of competing plasmid states whose subsequent dynamics may influence long-term memory persistence.

### Post-switching dynamics are governed by intracellular competition between incompatible plasmid states

The incomplete switching observed above implies that, immediately following induction, individual cells may contain a mixture of reporter plasmid variants differing in promoter orientation and therefore in transcriptional state. Because these variants share the same origin of replication, they can compete for the cellular replication and partitioning machinery ^23–28^. Such competition is well-established for plasmids belonging to the same incompatibility group, which generally cannot be stably maintained together within the same cell ^29–35,42^. We therefore asked whether intracellular competition between alternative plasmid states could account for the evolution of the heterogeneous population observed after switching.

We first focused on the first three days following switching (Fig. 2A–C). One day after induction, the majority of cells in both switching directions, RFP-to-GFP and GFP-to-RFP, were uncommitted, exhibiting both GFP and RFP fluorescence. Over the following days, the fraction of uncommitted cells progressively decreased, with cells resolving into either the GFP-positive or RFP-positive state. This progressive resolution suggests that cells initially containing both plasmid states undergo competition between incompatible plasmids, ultimately resulting in the dominance of one state within individual cells.

**Fig. 2.**
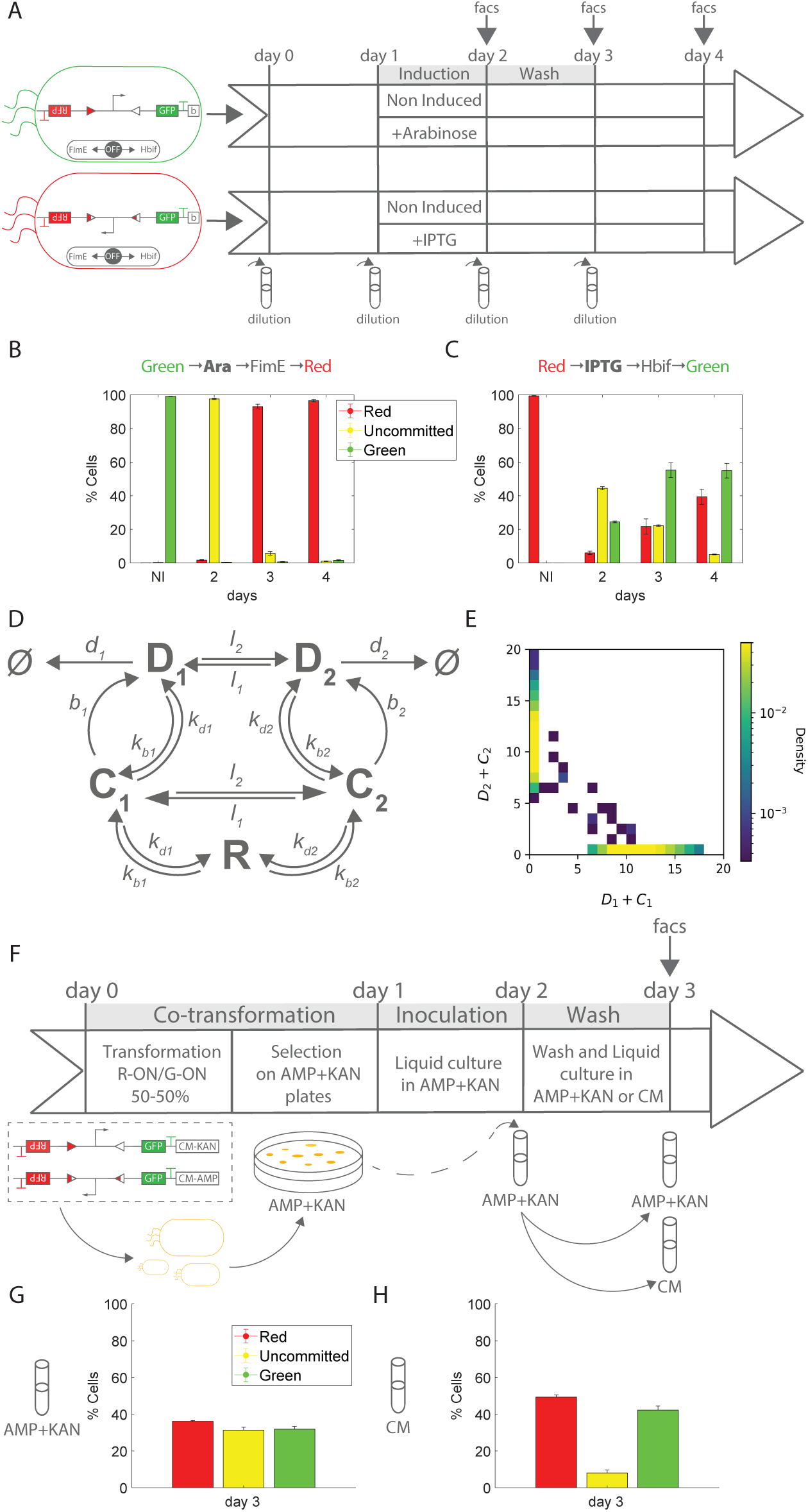
Competition between incompatible plasmids drives the dominance of one state over the other following the switch. (A) Schematic representation of the cell-switching treatment protocol. (B) Cells initialized in the Green-On state and switched to the Red-On state on days 2–4, together with the non-induced control (NI) analyzed on day 2. (C) Cells initialized in the Red-On state and switched to the Green-On state on days 2–4, together with the non-induced control (NI) analyzed on day 2. (D) Reaction diagram of the plasmid-competition model. (E) Stochastic model simulations of plasmid-competition dynamics showing the distribution of cells along the *D*_1_ + *C*_1_ and *D*_2_ + *C*_2_ axes. (F) Schematic representation of the experimental protocol used to investigate plasmid competition. (G) Cells co-transformed with inc^2^o^8^mpatible plasmids and maintained under ampicillin and kanamycin selection, analyzed on day 3. (H) Cells co-transformed with incompatible plasmids and maintained under chloramphenicol selection, analyzed on day 3.

To determine whether intracellular plasmid competition could account for these post-switching dynamics, we developed a stochastic model describing competition between the two incompatible plasmid variants (Fig. 2D), as reported in Methods. Based on the preceding results showing that trigger leakiness does not substantially affect the system over the experimental time scale, the model was constructed without including basal recombinase activity. Then, we initialized the model with equal copy numbers of the two plasmid states (D1 = D2 = 5), representing cells in an uncommitted state, and simulated their stochastic dynamics over time (Fig. 2E). The model predicts that stochastic competition between the two plasmid states drives individual cells toward one of two absorbing states, corresponding to the loss of one of the competing plasmid variants. Thus, the model recapitulates the experimentally observed transition from a population containing a large fraction of uncommitted cells to one dominated by either the GFP or RFP state.

Guided by these predictions, we designed an experiment to directly investigate competition between alternative plasmid states following a switch (Fig. 2F). To experimentally enforce the initial coexistence of the two states, we engineered reporter variants in the GFP-On and RFP-On configurations carrying distinct dual-antibiotic resistance markers. These plasmids were co-transformed into cells, generating populations in which both reporter plasmids shared the same origin of replication while conferring distinct antibiotic resistances, as described in the “Co-transformation experiments to study incompatibility competition” section of the Methods. Following co-transformation, cells were maintained either under dual-antibiotic selection, which favors retention of both plasmids, or under single-antibiotic selection, which allows competition between the incompatible plasmids to take place. Populations were monitored by flow cytometry (Fig. 2F).

Co-transformation of the two reporter plasmids resulted in a heterogeneous population comprising GFP-positive/RFP-positive, GFP-positive/RFP-negative, and GFP-negative/RFP-positive cells under dual-antibiotic selection (AMP + KAN; Fig. S2A). To determine whether this distribution was stable and reproducible, cells from the double-positive population were sorted and subsequently re-cultured (Fig. S2B). The following day, the same overall population distribution was recovered (Fig. S2C), indicating that the observed heterogeneity was not simply caused by transient measurement variability and was consistent with ongoing plasmid incompatibility dynamics. Interestingly, although cells were initially selected to retain both plasmids, a subset of cells displayed low or nearly undetectable fluorescence from one reporter. These cells may retain sufficient copies of the corresponding plasmid to confer antibiotic resistance while carrying insufficient copies, or expressing insufficient levels, of the reporter to produce detectable fluorescence.

Following transfer from dual-antibiotic to single antibiotic (chloramphenicol) selection, the fraction of GFP-positive/RFP-positive cells decreased sharply between days 2 and 3, with the population resolving predominantly into GFP-positive/RFP-negative or GFP-negative/RFP-positive states (Figs. 2G and H). This rapid resolution is consistent with ongoing competition between the two incompatible plasmids, whereby stochastic fluctuations in plasmid abundance progressively favor one plasmid over the other. The observation agrees with both the stochastic model prediction of transitions toward absorbing states and the progressive resolution observed following recombinase-mediated switching.

Notably, the transition occurred within a single overnight growth period, during which cells reached stationary phase. Thus, the rapid disappearance of the double-positive population is unlikely to be due to population fitness differences and instead supports intracellular plasmid competition as the primary mechanism underlying the observed early post-switch resolution.

These results establish that the heterogeneous genetic states generated during switching are not static. Rather, competing plasmid states undergo stochastic intracellular dynamics that rapidly resolve individual cells toward one of the two alternative memory states. We therefore next asked whether the processes governing this early resolution were sufficient to explain memory behavior over longer periods of population growth.

### Intercellular competition governs long-term state dynamics

The results above indicate that intracellular competition between incompatible plasmids can account for the rapid resolution of uncommitted cells into single-state populations following switching (Fig. 2). Once this intracellular resolution has occurred, however, cells carrying different memory states remain subject to population-level competition. If the two states impose different fitness costs, their relative abundance in the population may therefore continue to change even after the competing plasmid states have been resolved within individual cells. We therefore sought to distinguish the contribution of intracellular competition during the early post-switch phase from that of intercellular selection during long-term population growth.

We first monitored the system for ten days following switching (Fig. 3A–C). Based on the dynamics in Fig. 2, we considered the first three days to be primarily governed by intracellular competition between incompatible plasmids and therefore focused on the subsequent dynamics from day 4 onward. Following the GFP-to-RFP switch, the fraction of RFP-expressing cells remained relatively stable over this period (Fig. 3B). In contrast, following the RFP-to-GFP switch, the GFP-positive population reached approximately 60% before progressively decreasing, accompanied by an increase in the RFP-positive population (Fig. 3C). This divergence between the two switching directions suggests that, beyond the initial intracellular resolution of competing plasmid states, population-level selection favored the RFP-expressing state over longer time scales.

**Fig. 3.**
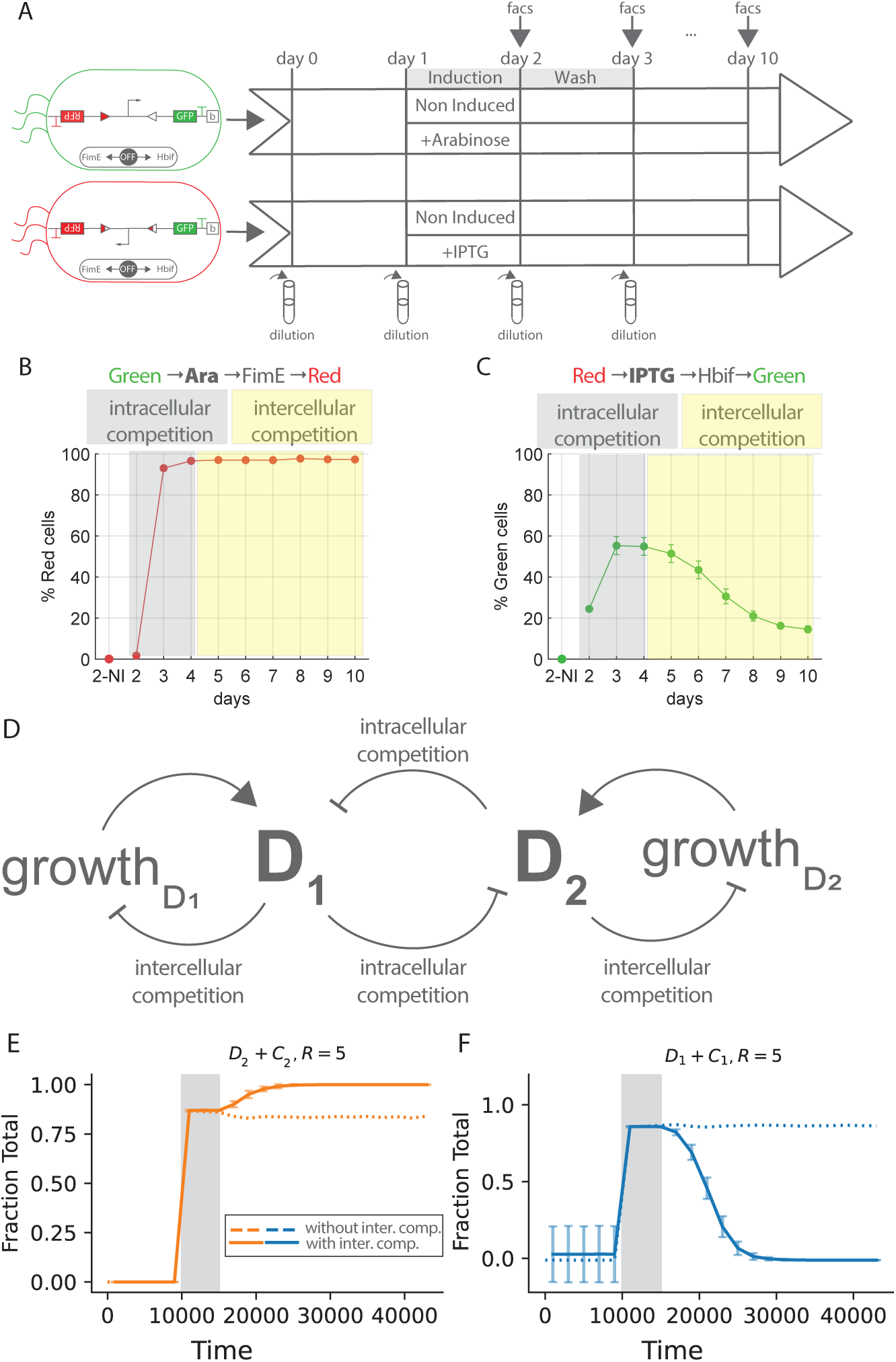
Intercellular dynamics govern the long-term maintenance of the system following the switch. (A) Schematic representation of the cell-switching treatment protocol. (B) % of red cells initialized in the Green-On state and switched to the Red-On state on days 2–10, together with the non-induced control (NI) analyzed on day 2. (C) % of green cells initialized in the Red-On state and switched to the Green-On state on days 2–10, together with the non-induced control (NI) analyzed on day 2. (D) Reaction diagram of the plasmid-competition model incorporating intercellular competition dynamics. (E) Stochastic model simulations of post-switch dynamics over time, with and without intercellular competition. Simulations induce the turning of D1 into D2. (F) Stochastic model simulations of post-switch dynamics over time, with and without intercellular competition. Simulations induce the turning of D2 into D1.

To test whether differences in cellular fitness could account for these long-term dynamics, we extended the stochastic model to incorporate cell replication and intercellular competition (Fig. 3D). Cell replication was introduced as an additional reaction with a propensity function defined as

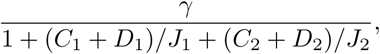

where *γ* represents the maximal growth rate and *J*_1_ and *J*_2_ determine the contribution of the corresponding plasmid-associated species to cellular fitness. Higher values of *J_x_*correspond to a lower fitness cost associated with the corresponding *D_x_* + *C_x_* species and therefore to higher relative cellular fitness. This formulation allows the model to capture differences in growth rates between cells carrying the two alternative plasmid states ^43^.

We then simulated the GFP-to-RFP (Fig. 3E) and RFP-to-GFP (Fig. 3F) switches using two model formulations, with or without population-level dynamics. For the GFP-to-RFP switch, both models reproduced the overall experimentally observed dynamics (Fig. 3B, E), although the model incorporating population-level dynamics more closely matched the experimental trajectory. In contrast, for the RFP-to-GFP switch, the model without population-level dynamics predicted stable maintenance of the GFP state, whereas incorporating differential cellular fitness reproduced the progressive loss of the GFP state observed experimentally (Fig. 3C, F). Thus, while intracellular plasmid competition captures the overall switching dynamics in both directions, inclusion of population-level fitness differences provides a more accurate description of the experimental trajectories and is necessary to reproduce the long-term loss of the GFP state.

Guided by these predictions, we next designed a control experiment to directly assess the contribution of intercellular competition. Reporter plasmids in the GFP-On and RFP-On states were introduced into separate bacterial populations, each together with the trigger plasmid, and the resulting populations were subsequently co-cultured (Fig. S3A), as described in the “Co-culturing experiments” section of the Methods. This experimental design prevents intracellular competition between the two reporter plasmids while allowing competition between the two resulting cell populations. The relative abundance of the two populations changed markedly between days 3, 7, and 13 (Fig. S3B). Over time, the co-cultures became increasingly enriched in RFP-expressing cells, indicating a fitness advantage of the RFP-expressing population under the conditions tested. These results show that population-level competition contributes substantially to the long-term dynamics of the system.

Taken together, these results reveal a temporal separation between two distinct processes governing the fate of the memory states. During the early post-switch phase, intracellular competition between incompatible plasmids drives the rapid resolution of heterogeneous cells into single-state populations (Fig. 2). On longer time scales, intercellular competition between cells carrying different states reshapes the population composition according to their relative fitness (Fig. 3).

### Plasmid copy number controls intracellular competition and long-term memory dynamics

Having identified intracellular plasmid competition and population-level fitness differences as distinct processes governing memory dynamics, we next asked whether these processes could be manipulated through circuit design. Because reporter plasmid copy number determines both the number of competing plasmid copies within individual cells and the level of reporter expression, we tested whether reducing copy number could reshape the post-switching dynamics at both stages.

We first examined the effect of plasmid copy number on intracellular state resolution using the stochastic model without the intercellular competition term (Fig. 4A). Lowering plasmid copy number accelerated the resolution of incompatible plasmid variants into the corresponding absorbing states, reducing the time during which the two states coexist within individual cells. The model therefore predicts faster resolution of mixed states at lower plasmid copy number.

**Fig. 4.**
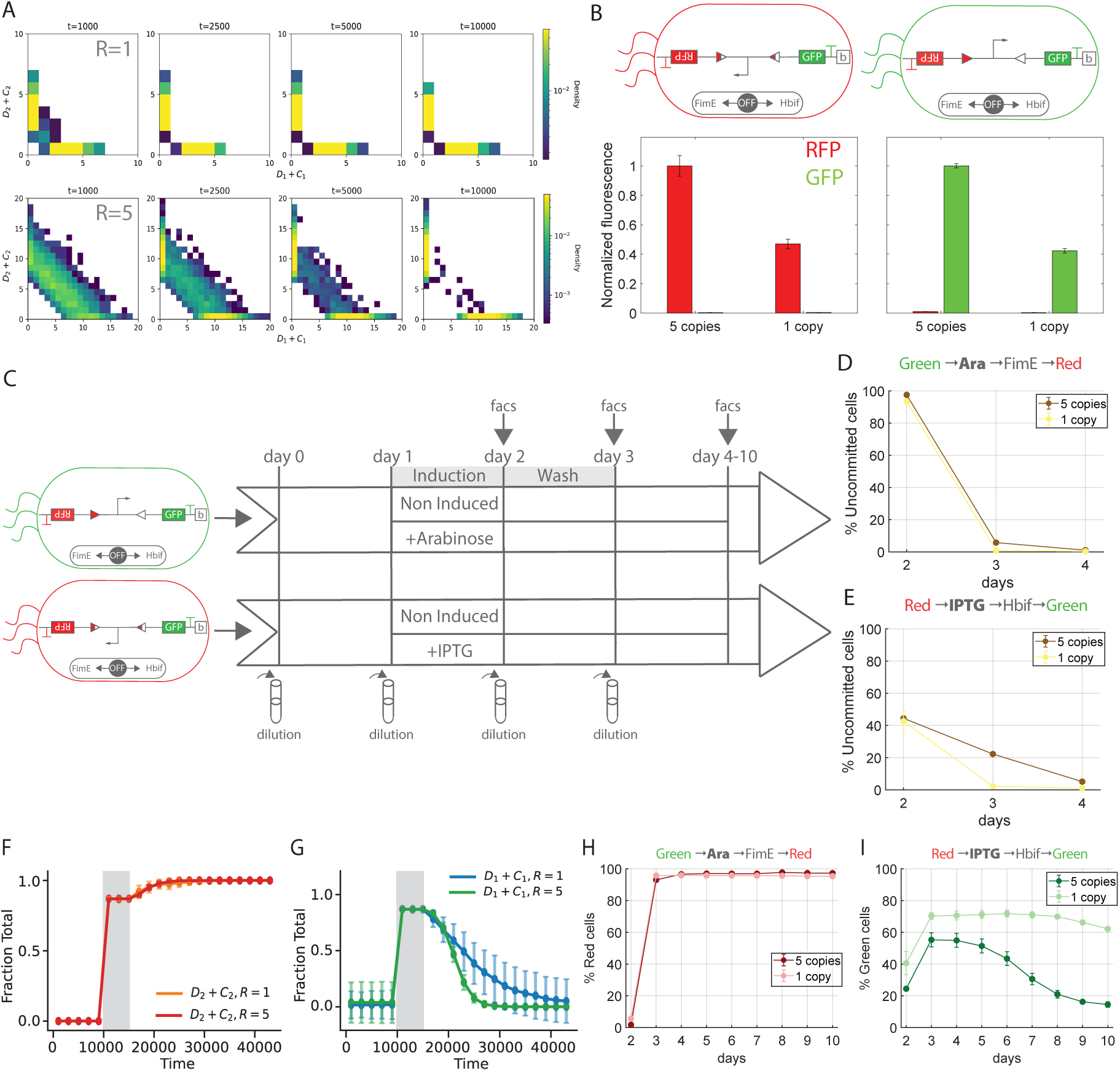
Plasmid copy number tunes intracellular state resolution and long-term memory persistence. (A) Stochastic model simulations of plasmid-competition dynamics showing the distribution of cells along the *D*_1_ +*C*_1_ and *D*_2_ + *C*_2_ axes as a function of time and plasmid copy number. (B) Normalized fluorescence as a function of plasmid copy number. Fluorescence was normalized to the five-copy system. (C) Schematic representation of the cell-switching treatment protocol. (D) Percentage of uncommitted cells initialized in the Green-On state and switched to the Red-On state on days 2–4. (E) Percentage of uncommitted cells initialized in the Red-On state and switched to the Green-On state on days 2–4. (F) Stochastic model simulations of post-switch dynamics over time as a function of plasmid copy number. Simulations induce the turning of D1 into D2. (G) Stochastic model simulations of post-switch dynamics over time as a function of plasmid copy number. Simulations induce the turning of D2 into D1. (H) Percentage of Red cells initialized in the Green-On state and switched to the Red-On state on days 2–10. (I) Percentage of Green cells initialized in the Red-On state and switched to the Green-On state on days 2–10.

To experimentally test this prediction, we generated a second reporter variant in which the reporter cassette was carried on a single-copy bacterial artificial chromosome (BAC) origin of replication ^44^. As expected, reducing the plasmid copy number resulted in a marked decrease in reporter fluorescence (Fig. 4B), confirming the lower expression associated with the single-copy configuration.

We then compared switching dynamics between the five-copy and single-copy systems (Fig. 4C). Following both the GFP-to-RFP and RFP-to-GFP transitions, we quantified the fraction of uncommitted cells over the subsequent three days (Fig. 4D,E). In both switching directions, the fraction of uncommitted cells decreased more rapidly in the single-copy system than in the five-copy system, consistent with the model prediction.

We further attempted to reproduce the co-transformation experiment described in Fig. 2F using two reporter plasmids carrying the same single-copy BAC origin. In contrast to the five-copy system, no colonies were recovered following co-transformation and selection for both plasmids. This result indicates that stable coexistence of the two reporter plasmids within the same cell is strongly disfavored in the single-copy configuration. The absence of stable double-plasmid transformants is consistent with the reduced capacity of the single-copy system to maintain competing reporter states within individual cells, although this result may also reflect intrinsic properties of BAC-based replication ^44,45^.

We next considered whether reduced copy number could also affect population-level competition. Because intercellular selection can arise from differences in cellular resource demand between the two reporter states (Fig. 3D), reducing reporter copy number is expected to decrease reporter-associated resource usage and thereby reduce fitness differences between RFP- and GFP-expressing cells. We therefore used the stochastic model incorporating intercellular competition to investigate whether long-term state persistence depends on plasmid copy number. Simulations of the GFP-to-RFP transition showed comparable dynamics across the two copy-number conditions (Fig. 4F). Simulations of the RFP-to-GFP transition showed that the long-term dynamics depended on plasmid copy number, with the single-copy system exhibiting a slower decline in the fraction of GFP-expressing cells than the five-copy system (Fig. 4G).

To experimentally test this prediction, we repeated the co-culture experiment using the single-copy reporter system (Fig. S3C). In contrast to the five-copy system, the relative abundance of RFP- and GFP-expressing populations remained substantially more balanced over time. Together with the reduced fluorescence observed for the single-copy system (Fig. 4B), these results indicate that lowering reporter copy number reduces the difference in cellular resource demand between the two states and consequently decreases their fitness difference.

We next assessed whether these changes translated into improved long-term state maintenance following switching. The GFP-to-RFP and RFP-to-GFP transitions were monitored over an extended time period in both the five-copy and single-copy systems. The two systems displayed comparable long-term behavior following the GFP-to-RFP transition (Fig. 4H). In contrast, the single-copy system showed a marked improvement following the RFP-to-GFP transition, exhibiting both higher switching efficiency and substantially improved maintenance of the GFP state compared with the five-copy system (Fig. 4I). These experimental results are consistent with the model predictions (Fig. 4F, G).

Taken together, these results demonstrate that plasmid copy number is a design parameter that influences memory dynamics at both intracellular and population levels. Lowering reporter copy number accelerates the resolution of competing plasmid states following switching while reducing differences in reporter-associated resource demand between alternative states. Thus, copy number provides a means of reshaping both the early resolution of heterogeneous states and the subsequent population-level dynamics, providing a route toward more persistent and balanced memory.

### Engineering relative state fitness to extend memory persistence

The reduction in reporter plasmid copy number improved both the resolution of competing states and long-term state maintenance. Hereafter, we refer to the single-copy configuration as S1-1 and to the five-copy configuration as S1-5. S1-1 and S1-5 maintained the RFP state for at least one week following the GFP-to-RFP transition, while the GFP state was maintained substantially better in S1-1 than in S1-5 between days 3 and 7 following the RFP-to-GFP transition, although a progressive loss of the GFP state was still observed thereafter (Fig. 4G,H). We therefore asked whether the residual fitness asymmetry in S1-1 could be further reduced to extend memory persistence.

To address this question, we modified the circuit architecture to directly tune the relative cellular resource demand of the two reporter states. As shown by the co-culture experiment (Fig. S3C), S1-1 exhibited substantially more balanced population dynamics than the five-copy system. However, a residual bias toward the RFP-expressing state was still apparent (Fig. 4G,H). We therefore hypothesized that further reducing the fitness advantage of the RFP-expressing population could improve maintenance of the GFP state, which was the state most affected by intercellular competition in S1-1 (Fig. 4H). This hypothesis follows the model framework introduced above (Fig. 3D), in which relative cellular fitness determines the long-term population composition following switching.

We therefore used the stochastic model including intercellular dynamics to simulate post-switch behavior as a function of relative cellular fitness while keeping the plasmid copy number fixed at one. Relative fitness was varied through the ratio of the *J*_1_ and *J*_2_ parameters, which determine the cellular cost associated with the two reporter states. The simulations confirmed that long-term state maintenance depends on relative cellular fitness (Fig. 5A). Reducing the fitness difference between the two states resulted in more balanced long-term maintenance, whereas increasing the fitness asymmetry promoted convergence toward the fitter state.

**Fig. 5.**
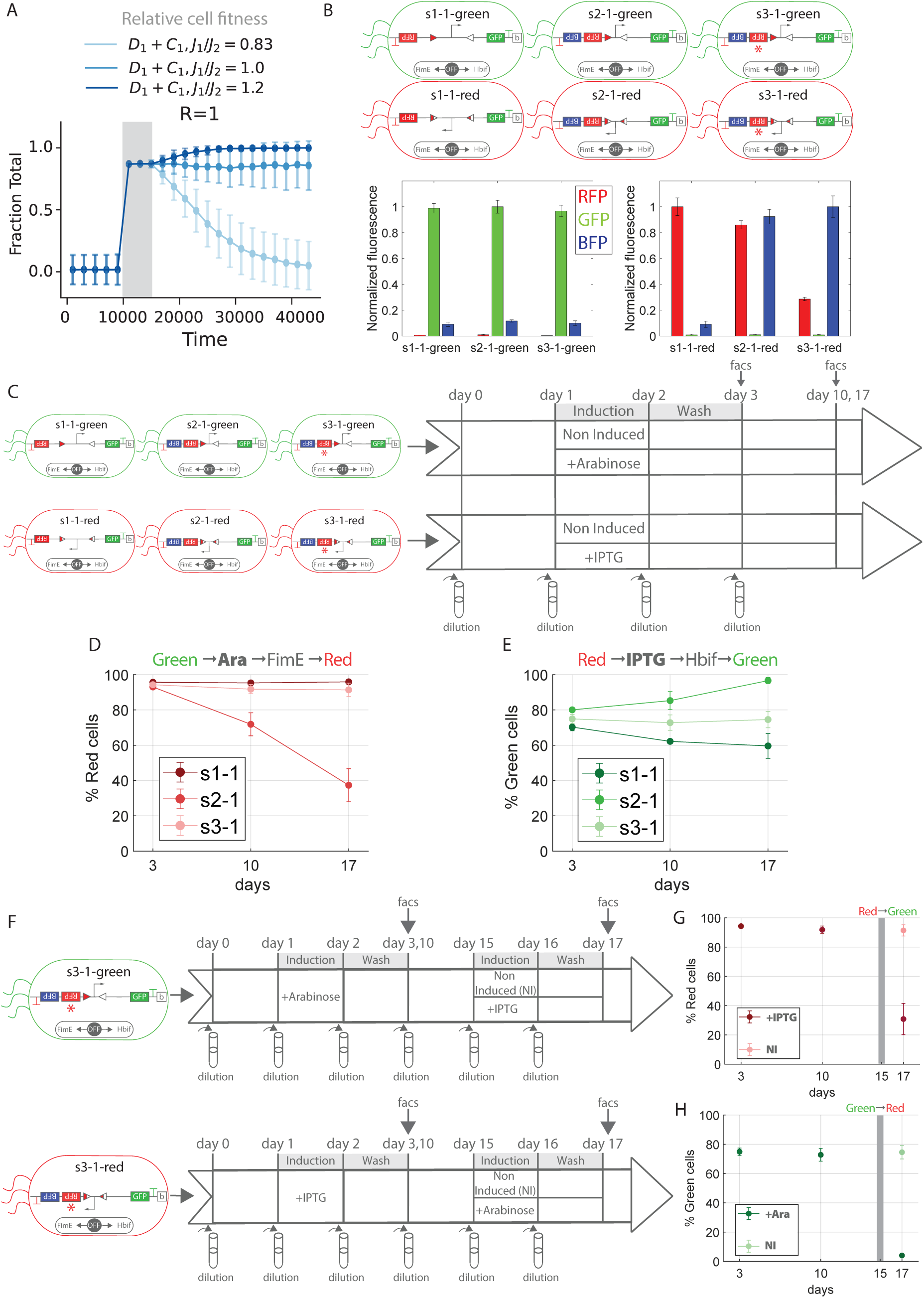
Tuning relative cellular fitness through circuit architecture improves long-term state maintenance. (A) Stochastic model simulations of post-switch dynamics over time for a plasmid copy number of 1 as a function of relative cell fitness. Simulations induce the turning of D2 into D1. (B) Normalized fluorescence for a plasmid copy number of 1 as a function of plasmid architecture. Fluorescence was normalized to the maximum fluorescence measured across the three systems for each fluorescence channel. (C) Schematic representation of the cell-switching treatment protocol. (D) Percentage of Red cells initialized in the Green-On state and switched to the Red-On state on days 3, 10, and 17. (E) Percentage of Green cells initialized in the Red-On state and switched to the Green-On state on days 3, 10, and 17. (F) Schematic representation of the cell-switching treatment protocol used to test rewritability of the optimized S3-1 system. Cells were initially switched on day 1 and maintained in the resulting state until day 15, when they were either subjected to a second switching treatment or left untreated as a control. Cell states were measured on days 3, 10, and 17. (G) Cells initialized in the Green-On state and switched to the Red-On state on day 1, followed by a second switching treatment back to the Green-On state on day 15. Cell states were measured on days 3, 10, and 17, together with the corresponding untreated control. (H) Cells initialized in the Red-On state and switched to the Green-On state on day 1, followed by a second switching treatment back to the Red-On state on day 15. Cell states were measured on days 3, 10, and 17, together with the corresponding untreated control.

We next sought to experimentally tune the relative cellular fitness of the two states in S1-1. Previous work suggests that loading on gene expression resources by the fluorescent reporter output can destabilize recombinase-based memory over extended periods ^13^. Rather than eliminating the loading associated with the stored state, we asked whether the relative loading of the two alternative states could be deliberately tuned to reduce their fitness asymmetry while retaining fluorescent outputs. Assuming that fluorescent protein production represents a major component of the cellular resource demand associated with the reporter, we hypothesized that increasing the resource demand of the RFP-expressing state could compensate for its residual fitness advantage and improve the stability of the GFP state.

To increase the resource demand specifically associated with the RFP state, we modified the red reporter cassette by introducing a second fluorescent protein, BFP, into the same transcriptional unit, generating a bicistronic reporter architecture. This new configuration, termed S2-1, resulted in an additional fluorescent output compared with the original S1-1 configuration (Fig. 5B).

The modified architecture substantially altered the population dynamics. In contrast to S1-1, which showed a balance between the RFP- and GFP-expressing states, S2-1 displayed a bias toward the GFP-expressing state in the co-culture experiment (Fig. S3D). Thus, increasing the resource demand of the RFP state was sufficient to flip the relative fitness of the two populations, demonstrating that the population-level bias could be quantitatively manipulated through circuit architecture.

We next asked whether this change in relative fitness translated into improved post-switch state maintenance. S1-1 and S2-1 were subjected to long-term switching experiments and monitored at days 3, 10, and 17 following induction (Fig. 5C). S2-1 showed improved switching and long-term maintenance following the RFP-to-GFP transition compared with S1-1 (Fig. 5E), consistent with the reduction of the original RFP fitness advantage. However, the increased resource demand of the red reporter overcompensated for the initial asymmetry, resulting in a new bias toward the GFP-expressing state. Consequently, the GFP-to-RFP transition exhibited reduced long-term stability (Fig. 5D). These results demonstrate that altering state-specific resource demand can tune both the magnitude and direction of the population-level bias, but that excessive compensation can flip the system toward the opposite state.

We therefore sought to generate an intermediate architecture that would balance the two states more effectively than either S1-1 or S2-1. Under the assumption that the majority of resource demand of gene expression is on translational resources ^46–49^, we lowered this demand by decreasing the ribosome binding site strength of the RFP gene ^49–51^. Starting from S2-1, we introduced a point mutation in the RFP ribosome-binding site to reduce its translation efficiency and consequently decrease RFP production (Fig. 5B). This intermediate configuration, termed S3-1, was designed to reduce the GFP-directed bias observed in S2-1 while retaining part of the increased resource demand introduced by the additional fluorescent protein.

The co-culture experiment confirmed that the modification reduced the bias toward the GFP-expressing state (Fig. S3E). We then evaluated the long-term switching performance of S3-1 using the same experimental design (Fig. 5C–E). Following the GFP-to-RFP transition, S3-1 displayed performance comparable to S1-1 (Fig. 5D). In contrast, following the RFP-to-GFP transition, S3-1 exhibited improved long-term state maintenance, with the GFP state remaining stable for up to two weeks (Fig. 5E).

Taken together, these results demonstrate that the relative fitness of alternative memory states can be deliberately tuned through circuit architecture. Starting from S1-1, increasing the resource demand associated with the RFP state shifted the population bias toward GFP, whereas partial reduction of this additional demand restored a more balanced system. Thus, the mechanistic framework established above can be converted into an engineering strategy: rather than simply maximizing switching efficiency, tuning the relative fitness of alternative states can reshape long-term population dynamics and extend the persistence of a reversible memory state.

### The optimized circuit retains rewritability after prolonged memory maintenance

We next assessed whether the optimized S3-1 circuit remained rewritable after prolonged memory maintenance. We followed the programmed states for two weeks after the initial switching event and then induced a second switching event in both directions (Fig. 5F). Specifically, cells maintained in the GFP-On state were switched to the RFP-On state (Fig. 5F, G), whereas cells maintained in the RFP-On state were switched to the GFP-On state (Fig. 5F, H). Both transitions remained functional after two weeks of memory maintenance, with switching efficiencies comparable to those observed at day 3 (Fig. 5G, H).

Thus, prolonged maintenance did not lock the circuit into its established state or impair access to the underlying recombination mechanism. S3-1 remained capable of rewriting its memory after two weeks. These results demonstrate that the extended persistence achieved through circuit engineering is compatible with continued reversibility, enabling a memory that is both long-lived and rewritable.

## Discussion

Our results show that the persistence of reversible genetic memory is not determined solely by the molecular mechanism that encodes the stored state. Instead, memory persistence emerges from a temporal hierarchy of processes operating within cells and across the growing population. In the system studied here, imperfect recombinase orthogonality generates heterogeneous genetic states during switching; these states are subsequently resolved through intracellular competition between incompatible plasmids, while longer-term population dynamics are governed by differences in the fitness of cells carrying alternative states. Thus, the mechanisms that establish a memory state are distinct from those that determine how that state persists.

This temporal separation provides a framework for understanding why switching efficiency alone is insufficient to predict memory persistence. Efficient recombination determines the initial distribution of states, but heterogeneous plasmid configurations can continue to evolve after the switching event, and the resulting populations remain subject to selection over many generations. Our results therefore distinguish two mechanistically different aspects of memory performance: the establishment of a desired state and the subsequent propagation of that state through a growing population. This distinction suggests that memory should be evaluated not only by how efficiently a state can be written, but also by how the resulting state evolves after inefficient writing.

More importantly, resolving these processes in time revealed distinct parameters through which memory dynamics could be engineered. Plasmid copy number provides one such design parameter because it affects both the number of competing plasmid copies within individual cells and the gene expression resource loading associated with the reporter. Previous work has similarly implicated gene copy number in the stability of recombinase-based reporters, with single-copy configurations showing more stable reporter states over extended cultivation ^52^. Our results extend this observation to reversible genetic memory by showing that copy number acts at different stages of the memory trajectory: reducing copy number accelerates the resolution of competing genetic states early after switching, while reducing reporter expression diminishes fitness differences between alternative states at later times. Thus, copy number can be used not simply to improve switching performance, but to reshape the subsequent dynamics of the memory state.

State-dependent cellular resource loading provides a complementary engineering parameter. Previous work showed that reducing the cellular resource loading associated with a stored state can improve the persistence of recombinase-based memory ^13^. In that study, persistence was improved by removing the fluorescent output associated with the stored state, thereby reducing its associated cellular resource loading but also eliminating the readout of the memory state. Our results suggest an alternative strategy that preserves this readout: rather than eliminating state-dependent cellular resource loading, the relative cellular resource loading of the alternative states can be tuned to reshape their fitness difference. By retaining fluorescent outputs while modulating their relative resource demand, we show that state-dependent cellular resource loading can be used to balance the long-term fitness of alternative memory states. More generally, the relevant design parameter for a reversible memory need not be the absolute cellular resource loading of either state, but the fitness difference between them. State-dependent cellular resource loading can therefore be transformed from a quantity to be minimized into a controllable parameter for engineering population-level memory dynamics.

Together, these findings suggest a broader design principle for genetic memory: rather than optimizing the switching event in isolation, memory persistence can be engineered by controlling the trajectory of the stored state after switching. In our system, this trajectory consists of at least two experimentally separable stages. Early dynamics are determined by the resolution of competing genetic configurations within individual cells, whereas later dynamics reflect the selective expansion of cells carrying alternative states. This decomposition makes it possible to target different stages with different interventions, controlling the extent and timescale of intracellular competition through plasmid architecture, and balancing population-level selection through state-specific resource demand. The resulting strategy converted mechanistic characterization into circuit engineering and extended memory persistence to approximately two weeks.

This approach also illustrates the limits of memory engineering based solely on circuit design. Even after reducing intracellular competition and balancing the relative fitness of the two states, residual fitness differences remain sufficient to bias the population over longer timescales. In a growing population, even small fitness differences can accumulate over many generations, progressively altering the population composition. Thus, while circuit engineering can substantially extend memory persistence, indefinite maintenance of a defined population state would require an additional mechanism capable of compensating for state loss or restoring the desired state over time. The relevant engineering objective is therefore not indefinite stability, but control over the timescale and trajectory of memory persistence.

Extending this timescale did not eliminate the functional reversibility of the circuit. The optimized S3-1 configuration remained capable of rewriting its stored state after two weeks of maintenance, showing that persistence and rewritability can be simultaneously achieved. At the same time, the reversible architecture introduces an additional constraint that is less pronounced in irreversible memory systems. Bidirectional switching requires two recombinases to act on the same memory substrate in opposite directions, creating a requirement for sufficiently orthogonal activities to avoid unintended recombination ^10,16,17^. In our system, incomplete orthogonality contributed to heterogeneous states during switching. By contrast, irreversible architectures based on a single recombinase and a defined pair of recognition sites avoid this particular constraint by writing the memory in a single direction ^13,17,52^. Thus, rewritable memories must balance access to both switching directions with sufficiently specific recombinase activity, adding an architectural consideration beyond long-term state stability.

Overall, our findings show that the long-term behavior of reversible genetic memory emerges from processes that unfold after the switching event. Rather than being defined solely by the efficiency with which a state is established, memory persistence can be shaped by controlling how that state evolves over time. This perspective provides a framework for designing genetic memories that combine prolonged persistence with direct readout and functional rewritability.

## Methods

### Cell lines

*E. coli* NEB5*α* cells (C2987H, New England Biolabs) were used in all cloning procedures. *E. coli* DH10b cells (EC0113, Thermo Fisher) were used in all experiments. *E. coli* DH10b do not have the *fimE/fimB* and *fim* structural genes and are not able to metabolize arabinose ^16^.

### Material

Cells were always grown in LB broth (LENNOX, 240230) using 13 mL VWR^®^ culture tubes with dual-position caps, polystyrene, clear tube (VWR, 60818-667). Chloramphenicol (34 *µ*g/mL, Sigma-Aldrich, C0378), ampicillin (100 *µ*g/mL, Sigma-Aldrich, A0166), and kanamycin (50 *µ*g/mL, Sigma-Aldrich, K1377) were supplemented when necessary ^16^. L-arabinose (1 mM, Sigma-Aldrich, A3256) and IPTG (1 mM, Sigma-Aldrich, I6758) were used as inducers ^16^. The pSwitch and pReporter plasmids ^16^ were kindly provided by Professor C. Voigt, Department of Biological Engineering, Massachusetts Institute of Technology (MIT), and served as the starting constructs for the engineering of the reporter system used in this study.

### Genetic circuits construction

The sensor plasmid modifications and assembly were based on Gibson assembly ^53^. DNA fragments to be assembled were amplified by PCR using Q5 High-Fidelity PCR Master Mix (NEB, M0492L), purified with gel electrophoresis and Zymoclean Gel DNA Recovery Kit (Zymo Research, D4002), quantified with the nanophotometer (Implen, P330), and assembled with Gibson Assembly Master Mix (NEB, E2611L). Assembled DNA was transformed into competent cells, NEB-5 competent E. coli (NEB, C2987H). Plasmid DNA was prepared by the plasmid miniprep-classic kit (Zymo Research, D4015). DNA sequencing used Plasmidsaurus DNA whole plasmid sequencing service.

### Flow Cytometry Measurements

The objective was to determine the distribution of RFP and GFP expression across individual cells in the culture using flow cytometry. All samples were analyzed on a BD FACSymphony A1 flow cytometer. Instrument templates were calibrated using untransformed bacterial cells for SSC-H vs SSC-A and SSC-W vs SSC-H plots to define the gating strategy. Positive control populations expressing either GFP or RFP were used to calibrate the fluorescence parameters: FITC-A (488 nm excitation, 530/30 nm emission filter) for green fluorescence, and PE-Texas Red-A (561 nm excitation, 610/20 nm emission filter) for red fluorescence. Each sample was diluted 1:500 in minimal medium and analyzed for 100,000 events to ensure statistical robustness.

### General cell culture and passaging conditions

The cell culture and passaging procedure was adapted from Fernandez-Rodriguez et al. ^16^ and was used as the standard procedure for all experiments described below. Briefly, DH10b cells carrying the indicated genetic circuits were inoculated daily in 950 *µ*L of LB medium supplemented with the appropriate antibiotics and incubated overnight at 37°C with shaking at 250 rpm. Unless otherwise specified, three biological replicates were used for each experimental condition.

On the first day, cells were inoculated from glycerol stocks or fresh colonies. On subsequent days, cultures were passaged by transferring 5 *µ*L of the previous day’s culture into fresh LB medium containing the appropriate antibiotics. When induction was required, the appropriate inducer was added according to the specific experimental protocol described below. For experiments involving long-term state maintenance, cells were washed twice in LB containing the appropriate antibiotics before being inoculated into fresh medium on the day following induction. This passaging procedure was repeated daily for the duration of each experiment. Flow cytometry was performed at the time points indicated for each experiment.

### Switching between states

Biological triplicates of each population were used to assess state switching. Cells were initialized according to the general cell culture and passaging procedure described above. On day 2, state switching was induced by adding 1 mM IPTG to induce the RFP-to-GFP transition or 1 mM L-arabinose to induce the GFP-to-RFP transition.

Following overnight induction, cells were washed twice in LB containing the appropriate antibiotics and reinoculated into fresh medium containing the same antibiotics but no inducer. Populations were subsequently monitored by flow cytometry (FACS) at the time points indicated in the main figures.

### Co-transformation experiments to study incompatibility competition

This protocol was developed to experimentally validate the modeling results describing competition between plasmids sharing the same origin of replication. The GFP-on and RFP-on reporter plasmids, both carrying the SC101 origin of replication, were engineered to contain dual antibiotic resistance markers (KAN-CM or AMP-CM). The two plasmids were co-transformed into cells in the following combination: GFP-on-KAN-CM + RFP-on-AMP-CM (Fig. 2F–H).

Co-transformed cells were plated under ampicillin and kanamycin selection (AMP + KAN) to ensure retention of both plasmids. The following day, three colonies from each condition were selected and inoculated into LB medium supplemented with AMP + KAN. After overnight growth, cells were washed and transferred to either LB + CM or LB + AMP + KAN and cultured for an additional day according to the general passaging procedure described above. Flow cytometry (FACS) analysis was performed on samples collected from overnight cultures.

### Co-culturing experiments

This protocol was developed to evaluate the contribution of population-level dynamics to the long-term maintenance of the cellular state following switching (Fig. S3). The GFP-on and RFP-on reporter plasmids, based on either the pSC101 or BAC origin of replication, were co-transformed into separate bacterial lines together with the trigger plasmid.

Transformed cells from each condition were plated under chloramphenicol and kanamycin selection (CM + KAN). The following day, three colonies per condition were selected and co-cultured in pairwise combinations. Each culture was inoculated at an initial OD_600_ of 0.01 in LB medium supplemented with CM + KAN. Cultures were maintained for 13 days using the general passaging procedure described above. Flow cytometry (FACS) analysis was performed on samples collected on days 3, 7, and 13.

### Plasmid competition model

The plasmid competition model of Figure 2D is comprised of the chemical reactions listed in Table 1. *D*_1_ and *D*_2_ represent two distinct plasmid constructs, *R* represents generic plasmid replication machinery, and *C*_1_ and *C*_2_ is the complex of *R* with *D*_1_ and *R* with *D*_2_, respectively. Reactions R_1_ and R_2_ capture the dilution of the plasmids due to cell division, R_3_ – R_6_ model the reversible binding of the plasmids with the DNA replication machinery *R*, reactions R_7_ and R_8_ capture plasmid duplication upon completion of DNA replication. Reactions R_9_ – R_12_ capture the flipping from one plasmid species to the other. R_13_ captures the rate of cell division for the population-level model.

**Table 1:** Plasmid competition reaction diagrams and rates.

| $R_i$ | Reaction | Prop. Function (Non-population model) | Prop. Function (Population model) |
| --- | --- | --- | --- |
| $R_1$ | $D_1 \xrightarrow{d_1} \emptyset$ | $a_1 = d_1 D_1$ | $a_1 = a_{13} D_1$ |
| $R_2$ | $D_2 \xrightarrow{d_2} \emptyset$ | $a_2 = d_2 D_2$ | $a_2 = a_{13} D_2$ |
| $R_3$ | $D_1 + R \xrightarrow{k_{b1}} C_1$ | $a_3 = \frac{k_{b1} D_1 R}{\Omega}$ | $a_3 = \frac{k_{b1} D_1 R}{\Omega}$ |
| $R_4$ | $C_1 \xrightarrow{k_{d1}} D_1 + R$ | $a_4 = k_{d1} C_1$ | $a_4 = k_{d1} C_1$ |
| $R_5$ | $D_2 + R \xrightarrow{k_{b2}} C_2$ | $a_5 = \frac{k_{b2} D_2 R}{\Omega}$ | $a_5 = \frac{k_{b2} D_2 R}{\Omega}$ |
| $R_6$ | $C_2 \xrightarrow{k_{d2}} D_2 + R$ | $a_6 = k_{d2} C_2$ | $a_6 = k_{d2} C_2$ |
| $R_7$ | $C_1 \xrightarrow{b_1} R + 2D_1$ | $a_7 = b_1 C_1$ | $a_7 = b_1 C_1$ |
| $R_8$ | $C_2 \xrightarrow{b_2} R + 2D_2$ | $a_8 = b_2 C_2$ | $a_8 = b_2 C_2$ |
| $R_9$ | $D_1 \xrightarrow{f_1 I_1 + f_{2,c} I_2 + f_{1,0}} D_2$ | $a_9 = D_1(f_1 I_1 + f_{2,c} I_2 + f_{1,0})$ | $a_9 = D_1(f_1 I_1 + f_{2,c} I_2 + f_{1,0})$ |
| $R_{10}$ | $D_2 \xrightarrow{f_2 I_2 + f_{1,c} I_1 + f_{2,0}} D_1$ | $a_{10} = D_2(f_2 I_2 + f_{1,c} I_1 + f_{2,0})$ | $a_{10} = D_2(f_2 I_2 + f_{1,c} I_1 + f_{2,0})$ |
| $R_{11}$ | $D_1 \xrightarrow{f_1 I_1 + f_{2,c} I_2 + f_{1,0}} D_2$ | $a_{11} = C_1(f_1 I_1 + f_{2,c} I_2 + f_{1,0})$ | $a_{11} = C_1(f_1 I_1 + f_{2,c} I_2 + f_{1,0})$ |
| $R_{12}$ | $D_2 \xrightarrow{f_2 I_2 + f_{1,c} I_1 + f_{2,0}} D_1$ | $a_{12} = C_2(f_2 I_2 + f_{1,c} I_1 + f_{2,0})$ | $a_{12} = C_2(f_2 I_2 + f_{1,c} I_1 + f_{2,0})$ |
| $R_{13}$ | Cell division | $a_{13} = 0$ | $a_{13} = \frac{\gamma}{1 + \frac{D_1 + C_1}{J_1} + \frac{D_2 + C_2}{J_2}}$ |

**Table 2:**
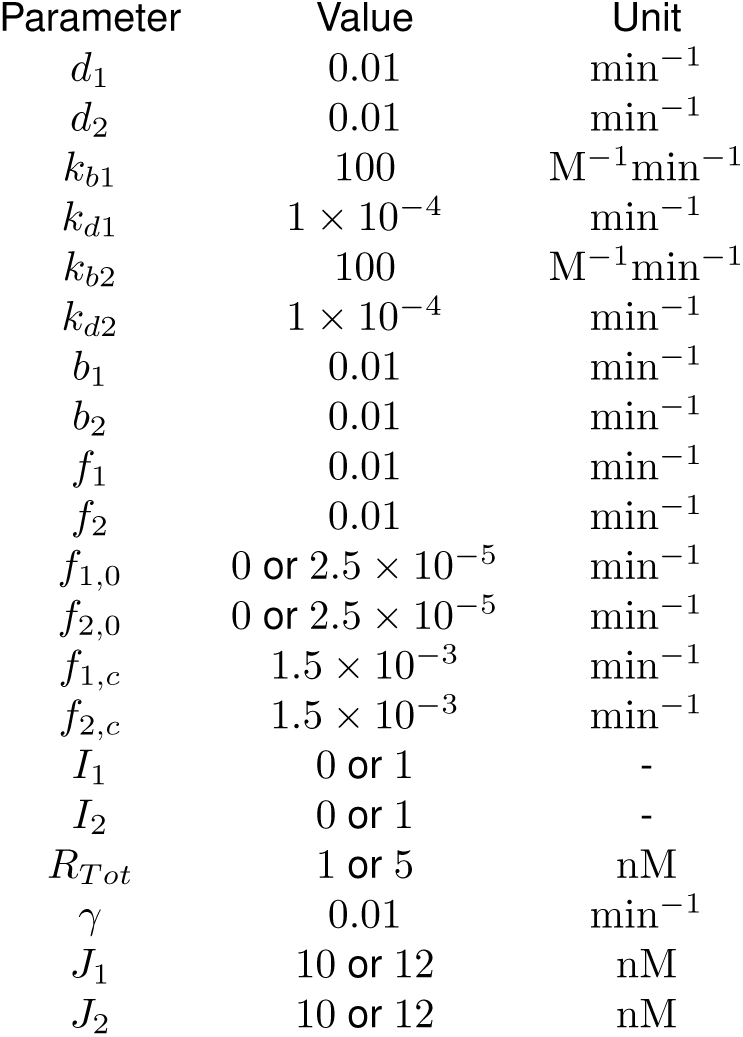
Plasmid competition parameters.

The model expands on the plasmid replication model proposed in Ruolo et al., in which the plasmid species *D*_1_ and *D*_2_ compete to be replicated by the same generic replication machinery *R* through the formation of complexes *C*_1_ and *C*_2_, which drive the synthesis of new plasmid DNA ^54^. Here we assume that both *D*_1_ and *C*_1_ can be converted into *D*_2_ and *C*_2_, respectively, and vice versa. Additionally, the flipping reactions accommodate for 1) induced flipping with rate constants *f*_1_ and *f*_2_, 2) leaky flipping where one species is spontaneously flipped to the other with rate constants *f*_1,0_ and *f*_2,0_, and 3) non-orthogonal flipping where induced flipping of one species can also flip the other species with rate constants *f*_1*,c*_ and *f*_2*,c*_. To select the rate constants for non-orthogonal flipping, we scanned an array of values and selected *f*_1*,c*_ = *f*_2*,c*_ = 1.5×10*^−^*^3^min*^−^*^1^ where the simulated behavior was comparable to experimental observations. The rate constants for leaky flipping were set to 0 min*^−^*^1^, follow with experimental observations.

Finally, we include the reaction R_13_ to model the rate of cell division for a population of bacteria with heterogeneous growth rates where the propensity function *a*_13_ is derived as in Hirsch et al. ^43^, and *a*_13_ is used to describe the dilution rate of plasmids for reactions R_1_ and R_2_ in the population-level model. *γ* = 0.01 min*^−^*^1^ is the basal growth rate constant and reflects a moderate population growth rate. *J*_1_ and *J*_2_ are the growth feedback parameters with values selected to reflect experimental observations.

The population-level model was implemented using the Next Family Method to simulate chemostat-type growth where *N* number of cells are simulated simultaneously, each as separate families containing reaction channels R_1_ – R_13_ ^55^. The growth rate is modified by Hirsch et al. ^43^. Upon firing of the cell division reaction R_13_, the daughter cell replaces another random cell in the population such that the population size is always constant.

---

**Fig. S1.**
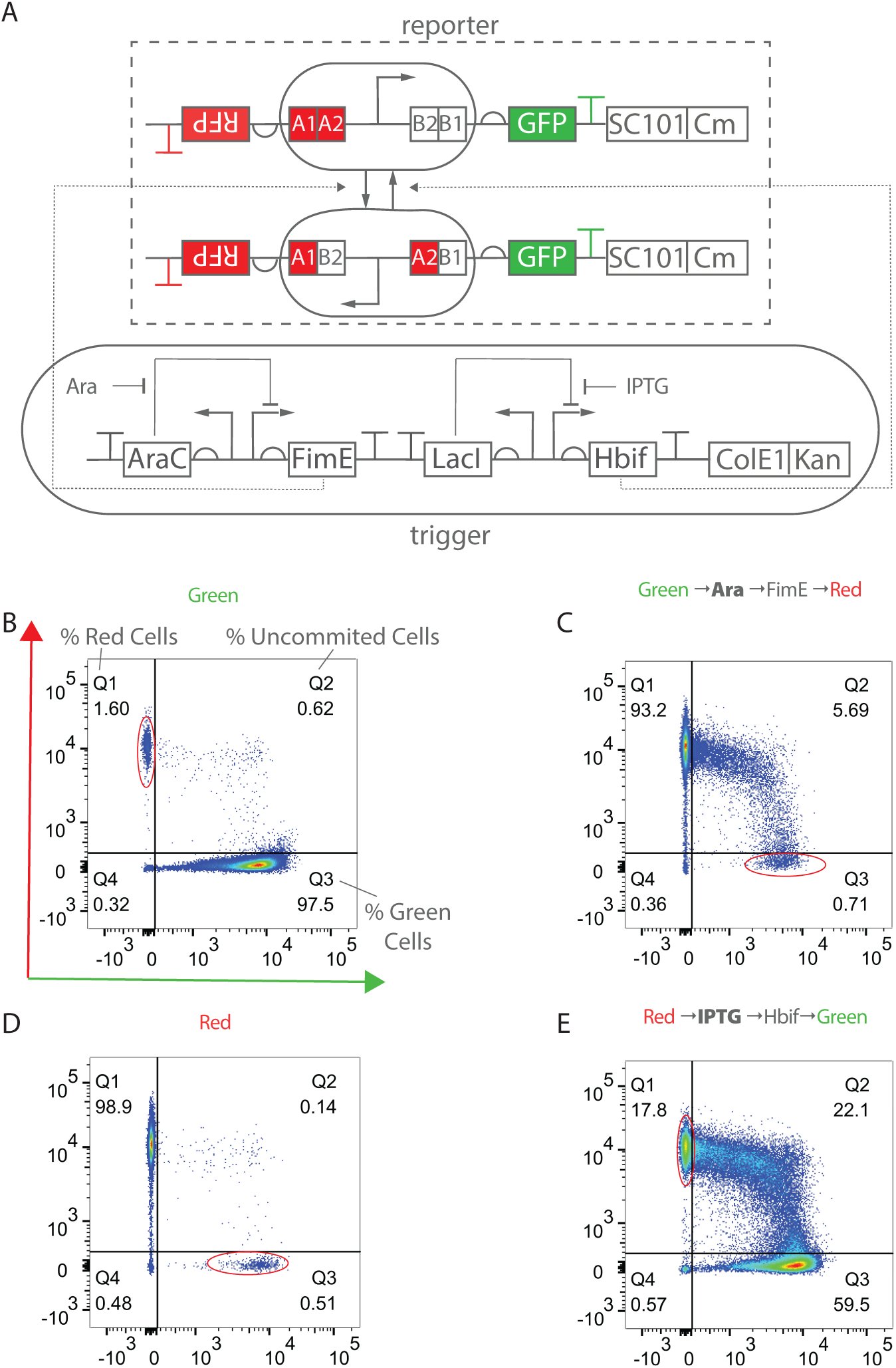
Recombinase-based bacterial switch: circuit architecture and flow cytometry characterization. (A) Architecture of the recombinase-based memory switch. The input module (pSwitch) contains IPTG- and arabinose-inducible promoters driving the expression of HbiF and FimE, respectively. The output module (pReporter) contains recombinase recognition sites flanking a constitutive promoter (Ptrc*) controlling RFP and GFP expression. Site-specific DNA inversion between the inverted repeats (IRL/IRR) reconfigures the recombination sites, reversing the promoter orientation and thereby toggling expression between RFP and GFP^16^. (B) Flow cytometry analysis of cells initialized in the Green-On state and analyzed on day 3. The percentages of Red, uncommitted, and Green cells are reported as indicated. Red circles indicate the effect of recombinase leakiness and non-orthogonality. (C) Flow cytometry analysis of cells initialized in the Green-On state and switched to the Red-On state on day 3. (D) Flow cytometry analysis of cells initialized in the Red-On state and analyzed on day 3. (E) Flow cytometry analysis of cells initialized in the Red-On state and switched to the Green-On state on day 3.

**Fig. S2.**
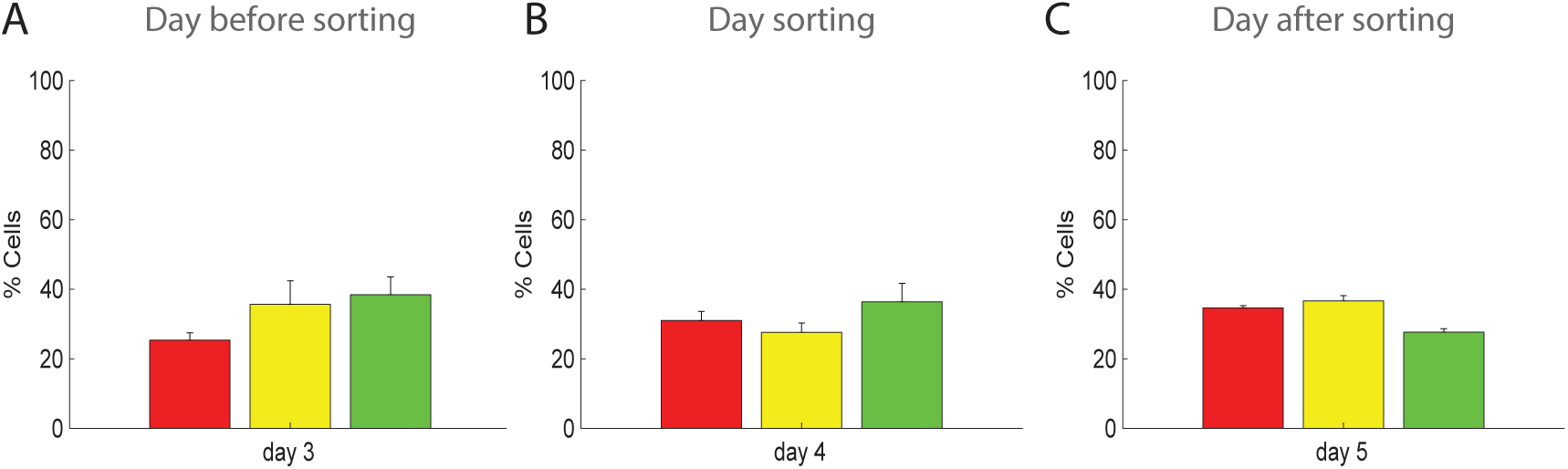
Sorting control confirms plasmid competition under dual-antibiotic selection. Populations co-transformed as shown in Fig. 2 were sorted based on red- and green-positive fluorescence and subsequently re-cultured to confirm the population distribution observed by flow cytometry. (A) Flow cytometry analysis performed on the day before sorting. (B) Flow cytometry analysis performed on the day of sorting. (C) Flow cytometry analysis performed on the day after sorting.

**Fig. S3.**
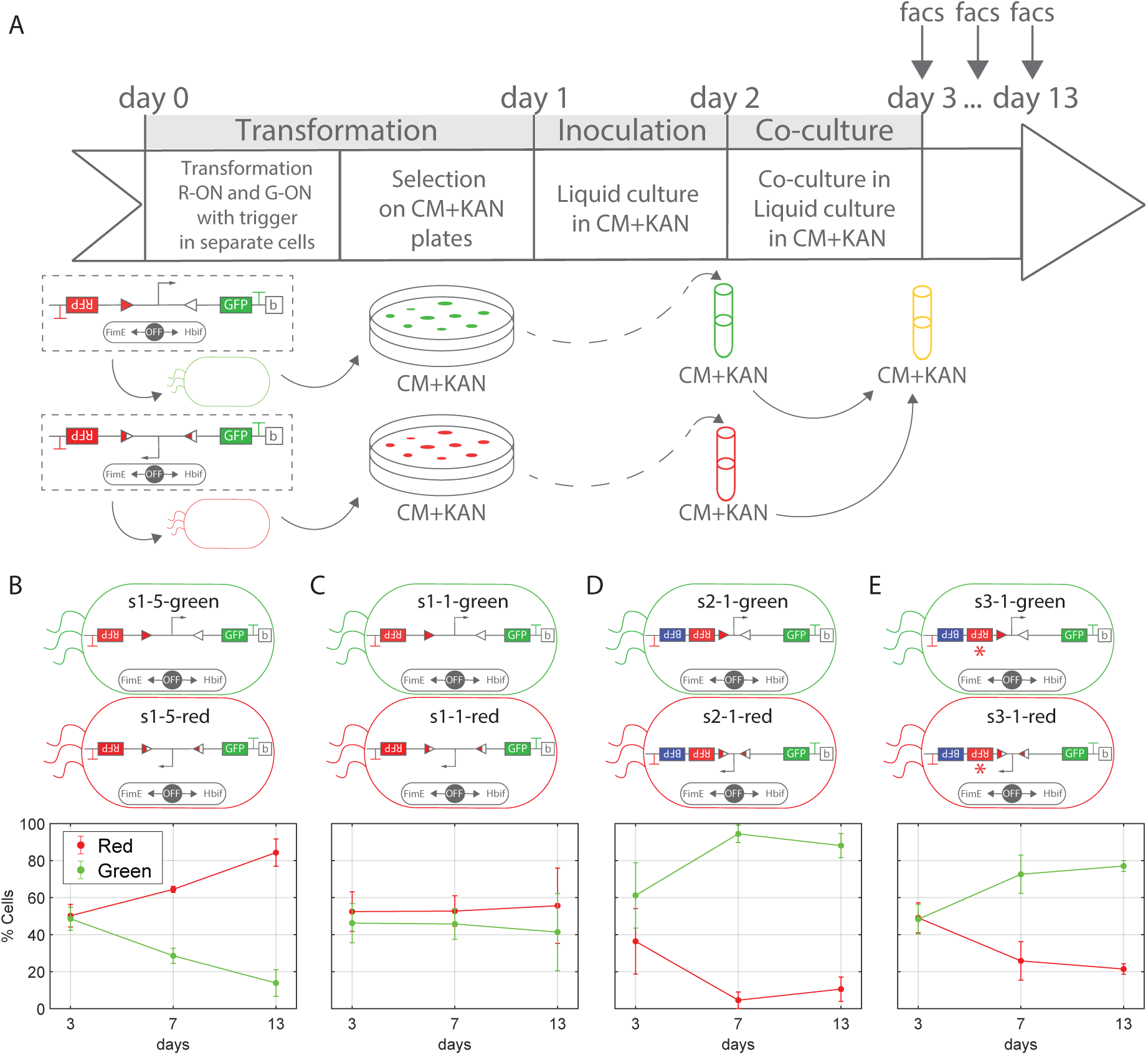
Engineering translational resource loading enables biasing of population dynamics. (A) Schematic representation of the cell treatment protocol. Populations carrying reporter constructs initialized in either the RFP or GFP state and transformed with the trigger plasmid were co-cultured to assess the contribution of population dynamics across different plasmid architectures. (B) Population distributions measured by flow cytometry on days 3, 7, and 13 for System 1 at a 5-copy plasmid configuration. (C) Population distributions measured by flow cytometry on days 3, 7, and 13 for System 1 at a 1-copy plasmid configuration. (D) Population distributions measured by flow cytometry on days 3, 7, and 13 for System 2 at a 1-copy plasmid configuration. (E) Population distributions measured by flow cytometry on days 3, 7, and 13 for System 3 at a 1-copy plasmid configuration.

## References

[1] S. Huang et al., “Bifurcation dynamics in lineage-commitment in bipotent progenitor cells,” Developmental Biology, vol. 305, no. 2, pp. 695–713, 2007.

[2] T. S. Gardner et al., “Construction of a genetic toggle switch in escherichia coli,” Nature, vol. 403, pp. 339–342, 2000.

[3] N. Dalchau et al., “Computing with biological switches and clocks,” Natural Computing, vol. 17, pp. 761–779, 2018.

[4] P. Siuti et al., “Synthetic circuits integrating logic and memory in living cells,” Nature Biotechnology, vol. 31, pp. 448–452, 2013.

[5] O. Purcell et al., “Synthetic analog and digital circuits for cellular computation and memory,” Current Opinion in Biotechnology, vol. 29, pp. 146–155, 2014.

[6] N. Roquet, A. P. Soleimany, A. C. Ferris, S. Aaronson, and T. K. Lu, “Synthetic recombinase-based state machines in living cells,” Science, vol. 353, no. 6297, p. aad8559, 2016.

[7] P.-F. Xia et al., “Synthetic genetic circuits for programmable biological functionalities,” Biotechnology Advances, vol. 37, p. 107393, 2019.

[8] D. T. Riglar et al., “Engineered bacteria can function in the mammalian gut long-term as live diagnostics of inflammation,” Nature Biotechnology, vol. 35, pp. 653–658, 2017.

[9] J. W. Lee, C. T. Chan, S. Slomovic, and J. J. Collins, “Next-generation biocontainment systems for engineered organisms,” Nature chemical biology, vol. 14, no. 6, pp. 530–537, 2018.

[10] J. Bonnet et al., “Rewritable digital data storage in live cells via engineered control of recombination directionality,” Proceedings of the National Academy of Sciences, vol. 109, no. 23, pp. 8884–8889, 2012.

[11] B. An, T.-C. Tang, Q. Zhang, T. Wang, Y. Wang, K. Gan, K. Liu, D. L. Zhang, Y. Liu, Y. K. Pan et al., “Synthetic circuits for cell ratio control,” Nature, vol. 653, no. 8114, pp. 587–598, 2026.

[12] D. Graf, L. Laistner, V. Klingel, N. E. Radde, S. Weirich, and A. Jeltsch, “Reversible switching and stability of the epigenetic memory system in bacteria,” The FEBS Journal, vol. 290, no. 8, pp. 2115–2126, 2023.

[13] P. B. Kalvapalle et al., “Long-duration environmental biosensing by recording analyte detection in dna using recombinase memory,” Applied and Environmental Microbiology, vol. 90, no. 4, 2024.

[14] U. Kwon et al., “Design of a long-term memory genetic toggle switch inspired by chromatin modification circuits,” in IEEE 61st Conference on Decision and Control (CDC), 2022.

[15] N. D. Grindley, K. L. Whiteson, and P. A. Rice, “Mechanisms of site-specific recombination,” Annu. Rev. Biochem., vol. 75, no. 1, pp. 567–605, 2006.

[16] J. Fernandez-Rodriguez et al., “Memory and combinatorial logic based on dna inversions: dynamics and evolutionary stability,” ACS Synthetic Biology, vol. 4, no. 12, pp. 1361–1372, 2015.

[17] L. Yang et al., “Permanent genetic memory with >1-byte capacity,” Nature Methods, vol. 11, no. 12, pp. 1261–1266, 2014.

[18] J. M. Abraham, C. S. Freitag, J. R. Clements, and B. I. Eisenstein, “An invertible element of dna controls phase variation of type 1 fimbriae of escherichia coli.” Proceedings of the National Academy of Sciences, vol. 82, no. 17, pp. 5724–5727, 1985.

[19] J. Dworkin and M. J. Blaser, “Generation of campylobacter fetus s-layer protein diversity utilizes a single promoter on an invertible dna segment,” Molecular microbiology, vol. 19, no. 6, pp. 1241–1253, 1996.

[20] P. Klemm, B. J. Jørgensen, I. van Die, H. de Ree, and H. Bergmans, “The fim genes responsible for synthesis of type 1 fimbriae in escherichia coli, cloning and genetic organization,” Molecular and General Genetics MGG, vol. 199, no. 3, pp. 410–414, 1985.

[21] K. G. Weinacht, H. Roche, C. M. Krinos, M. J. Coyne, J. Parkhill, and L. E. Comstock, “Tyrosine site-specific recombinases mediate dna inversions affecting the expression of outer surface proteins of bacteroides fragilis,” Molecular Microbiology, vol. 53, no. 5, pp. 1319–1330, 2004.

[22] B. D. Huang, D. Kim, Y. Yu, and C. J. Wilson, “Engineering intelligent chassis cells via recombinase-based memory circuits,” Nature Communications, vol. 15, no. 1, p. 2418, 2024.

[23] G. Selzer et al., “The origin of replication of plasmid p15a and comparative studies on the nucleotide sequences around the origin of related plasmids,” Cell, vol. 32, no. 1, pp. 119–129, 1983.

[24] I. Freudenau et al., “Cole1-plasmid production in escherichia coli: mathematical simulation and experimental validation,” Frontiers in Bioengineering and Biotechnology, vol. 3, p. 127, 2015.

[25] M. S. Standley et al., “Genetic control of cole1 plasmid stability that is independent of plasmid copy number regulation,” Current Genetics, vol. 65, no. 1, pp. 179–192, 2019.

[26] V. Brendel et al., “Quantitative model of cole1 plasmid copy number control,” Journal of Molecular Biology, vol. 229, no. 4, pp. 860–872, 1993.

[27] J. D. Keasling et al., “Cole1 plasmid replication: a simple kinetic description from a structured model,” Journal of Theoretical Biology, vol. 141, no. 4, pp. 447–461, 1989.

[28] J. Paulsson et al., “Requirements for rapid plasmid cole1 copy number adjustments: a mathematical model of inhibition modes and rna turnover rates,” Plasmid, vol. 39, no. 3, pp. 215–234, 1998.

[29] F. Rossine et al., “Intracellular competition shapes plasmid population dynamics.” Science, vol. 6779, no. 390, p. eadx0665, 2025.

[30] R. P. Novick, “Plasmid incompatibility,” Microbiological Reviews, vol. 51, no. 4, pp. 381–395, 1987.

[31] G. del Solar et al., “Replication and control of circular bacterial plasmids,” Microbiology and Molecular Biology Reviews, vol. 62, no. 2, pp. 434–464, 1998.

[32] R. Diaz et al., “Imaging centromere-based incompatibilities: insights into the mechanism of incompatibility mediated by low-copy number plasmids,” Plasmid, vol. 80, pp. 54–62, 2015.

[33] G. Ebersbach et al., “Partition-associated incompatibility caused by random assortment of pure plasmid clusters,” Molecular Microbiology, vol. 56, no. 6, pp. 1430–1440, 2005.

[34] M. A. Schumacher, “Bacterial plasmid partition machinery: a minimalist approach to survival,” Current Opinion in Structural Biology, vol. 22, no. 1, pp. 72–79, 2012.

[35] N. van der Hoeven, “Coexistence of incompatible plasmids in a bacterial population living under a feast and famine regime,” Journal of Mathematical Biology, vol. 24, no. 3, pp. 313–325, 1986.

[36] S. Kumar, A. Lezia, and J. Hasty, “Engineering plasmid copy number heterogeneity for dynamic microbial adaptation,” Nature microbiology, vol. 9, no. 8, pp. 2173–2184, 2024.

[37] N. F. Hülter, T. Wein, J. Effe, A. Garoña, and T. Dagan, “Intracellular competitions reveal determinants of plasmid evolutionary success,” Frontiers in microbiology, vol. 11, p. 2062, 2020.

[38] S. Bedhomme, D. Perez Pantoja, and I. G. Bravo, “Plasmid and clonal interference during post horizontal gene transfer evolution,” Molecular Ecology, vol. 26, no. 7, pp. 1832–1847, 2017.

[39] K. Yamaguchi and M. Yamaguchi, “The replication origin of psc101: the nucleotide sequence and replication functions of the ori region,” Gene, vol. 29, no. 1–2, pp. 211–219, 1984.

[40] A. A. K. Nielsen et al., “Multi-input crispr/cas genetic circuits that interface host regulatory networks,” Molecular Systems Biology, vol. 10, no. 11, p. 763, 2014.

[41] Y. Qian et al., “Realizing integral control in living cells: how to overcome leaky integration due to dilution?” Journal of the Royal Society Interface, vol. 14, no. 131, p. 20160833, 2018.

[42] M. Shintani et al., “Genomics of microbial plasmids: classification and identification based on replication and transfer systems and host taxonomy,” Frontiers in Microbiology, vol. 6, p. 242, 2015.

43. D. Hirsch and D. Del Vecchio, “Differential equation model for the population-level dynamics of a toggle switch with growth-feedback,” in 2022 IEEE 61st Conference on Decision and Control (CDC). IEEE, 2022, pp. 3207–3212.

[44] H. Shizuya, B. Birren, U. Kim, V. Mancino, T. Slepak, Y. Tachiiri, and M. Simon, “Cloning and stable maintenance of 300-kilobase-pair fragments of human dna in *escherichia coli* using an f-factor-based vector,” Proceedings of the National Academy of Sciences, vol. 89, no. 18, pp. 8794–8797, 1992.

[45] D. R. Williams and C. M. Thomas, “Active partitioning of bacterial plasmids,” Microbiology, vol. 138, no. 1, pp. 1–16, 1992.

[46] M. Scott, C. W. Gunderson, E. S. Mateescu, Z. Zhang, and T. Hwa, “Interdependence of cell growth and gene expression: origins and consequences,” Science, vol. 330, no. 6007, pp. 1099–1102, 2010.

[47] R. Balakrishnan et al., “Suboptimal resource allocation in changing environments constrains response and growth in bacteria,” Molecular Systems Biology, vol. 17, no. 12, p. MSB202110597, 2021.

[48] X. Dai et al., “Reduction of translating ribosomes enables Escherichia coli to maintain elongation rates during slow growth,” Nature Microbiology, vol. 2, no. 2, p. 16231, 2016.

[49] A. Gyorgy, J. I. Jiménez, J. Yazbek, H.-H. Huang, H. Chung, R. Weiss, and D. Del Vecchio, “Isocost lines describe the cellular economy of genetic circuits,” Biophysical journal, vol. 109, no. 3, pp. 639–646, 2015.

[50] C. Barajas et al., “Feedforward growth rate control mitigates gene activation burden,” Nature Communications, vol. 13, no. 1, p. 7054, 2022.

[51] Y. Qian et al., “Resource competition shapes the response of genetic circuits,” ACS Synthetic Biology, vol. 6, no. 7, pp. 1263–1272, 2017.

[52] E. Garabello, H. Yoon, M. C. Reid, and A. Giometto, “Tunable low-rate genomic recombination with cre-lox in Escherichia coli: a versatile tool for anoxic environmental biosensing and synthetic biology,” Applied and Environmental Microbiology, vol. 92, no. 4, p. e0176825, 2026.

[53] D. G. Gibson, L. Young, R.-Y. Chuang, J. C. Venter, C. A. Hutchison III, and H. O. Smith, “Enzymatic assembly of dna molecules up to several hundred kilobases,” Nature methods, vol. 6, no. 5, pp. 343–345, 2009.

54. I. Ruolo, E. Lu, and D. Del Vecchio, “Stochastic analysis of plasmids competition in a bacterial switch,” in 2026 American Control Conference (ACC). IEEE, 2026, pp. 1904–1909.

[55] A. Roy and S. Klumpp, “Simulating genetic circuits in bacterial populations with growth heterogeneity,” Biophysical Journal, vol. 114, no. 2, pp. 484–492, 2018.

